# Importance of functional diversity in benthic remineralization: a new perspective through the lens of Nares Strait, a key Arctic gateway

**DOI:** 10.64898/2026.03.11.710703

**Authors:** Thibaud Combaz, Bodil A. Bluhm, Ursula Witte, Philippe Archambault

## Abstract

Benthic remineralization of organic matter is key to carbon and nutrient cycling, influencing both long-term carbon storage in the sediments and the release of nutrients that support primary production in the water column. With its multiple forms and ages of sea ice, Nares Strait in the Canadian Arctic offers a unique opportunity to address the knowledge gap of variability of benthic remineralization rates along a natural sea ice gradient. Here, we incubated sediment cores in different locations in Nares Strait characterised by different sea ice conditions ranging from first-year ice to multi-year ice, to measure oxygen and nutrient fluxes. To identify potential drivers, we measured environmental variables, identified macrofauna and calculated a suite of taxonomic and functional diversity indices. Our analyses showed that benthic fluxes varied significantly between the northern and southern regions of Nares Strait. The presence of deposit feeders and sea ice cover (number of days since ice-free) were the main drivers in benthic fluxes, explaining 22.6% and 13.9% of the benthic flux variation, respectively. Overall, functional diversity was a better predictor of benthic fluxes than taxonomic diversity, indicating its primary importance in controlling benthic ecosystems functioning. Our results reveal that, from a benthic biogeochemical point of view, Nares Strait seems to be dissected into two main sub-regions: (i) a permanently and highly sea ice-covered area north of Kennedy Channel, resembling deeper regions of the Arctic Ocean and (ii) a seasonally ice-covered area between the North Water Polynya and Kane Basin, where benthic fluxes values are equivalent to those reported in similar continental Arctic shelves. Consequently, the rapid functional shifts resulting from the ongoing decline in sea ice could enhance benthic remineralisation rates if deposit feeder were to become dominant in certain areas, reducing the role of the region and by extension, the Arctic, as a carbon sink.

## Introduction

Ocean sediments represent Earth’s largest surface for carbon storage, yet most models ignore or minimally consider the diverse seafloor habitats [1]. It is becoming increasingly clear that benthic communities provide a key ecosystem service by effectively removing carbon from the atmosphere and either burying or recycling it in sediments [2]. Specifically, organic matter remineralization refers to the breakdown of organic matter into its simplest inorganic forms, such as nutrients and carbon. This process is crucial for ecosystems because it recycles organic matter, making it available again for reuse by primary producers. While remineralisation occurs throughout the entire water column, benthic (seafloor) remineralization ultimately determines the ocean’s long-term carbon storage capacity, as it controls whether carbon is released back into the water column or is buried in sediments [3,4]. Yet, the drivers of benthic remineralization are still poorly understood on regional scales. Globally, environmental conditions such as organic matter availability, temperature, salinity and depth interact and influence biodiversity, which in turn influences remineralization rates [5–8]. the interplay of these factors produces complex biodiversity and remineralization patterns that can vary on regional and global scales [9].

Historically, research has quantified biodiversity using “classical” taxonomic metrics such as taxa richness, abundance or biomass. However, several recent studies have pointed out that rather functional diversity within a community explains the magnitude of ecosystem processes [10–13]. Functional diversity refers to the set of physiological, morphological and behavioural characteristics of species that determine the ecological roles of organisms and how they interact with their environment [14,15]. Recently, much work has been done to develop indices that capture the functional diversity in ecosystems [16–18].

In the Arctic, the availability of labile organic matter is one of the most important parameter in driving remineralization rates [19]. Sea ice plays an important role in primary production by reducing the amount of light available, thereby affecting the timing of both the sympagic and pelagic spring blooms [20,21]. Food supply to the benthos in the Arctic is highly seasonal and is tightly linked to sea ice dynamics. As a result of climate change and consequent decrease in sea ice extent and cover, the Arctic-wide open water area (i.e., ice-free) has already increased by 27% between 1998 and 2018 [22] and current climate models suggest these trends will continue [23,24]. Total annual primary production is predicted to increase in many areas [22,25]. These changes are likely to impact the structure and the functions of Arctic benthic ecosystems, with possible implications for carbo burial and nutrient recycling [26].

Our study focuses on Nares Strait, located between Canada and Greenland. This region, one of the least explored regions of the world ocean, is unique because it constitutes an important connection between the Arctic Ocean and the Atlantic Ocean, via Baffin Bay. It acts as a transition zone between the Lincoln Sea in the north, where most of the remaining multi-year ice in the Arctic Ocean is found, and the highly productive *Pikialasorsuaq region* or North Water Polynya (NOW) in the south, one of the most productive polynyas in the Arctic. Nares Strait is also part of the Last Ice Area, which is expected to persist as the central Arctic Ocean becomes seasonally ice-free in a few decades [27]. With its multiple forms and ages of sea ice, Nares Strait acts as a sentinel of the processes affecting the Arctic Ocean and, therefore, as a crucial in situ “laboratory” for studying the evolution of marine ecosystems in a changing environment [28]. The objectives of this study are thus i) to quantify the oxygen uptake and nutrient fluxes of the benthic communities in the different subregions, ii) to evaluate potential environmental drivers of benthic remineralization, and iii) to assess the respective influence of taxonomic and functional diversity on this process.

## Materials and methods

### Study site

Nares Strait is a narrow passage along which sea ice and freshwater leave the Arctic Ocean into sub-Arctic waters next to the outflow in western Fram Strait. The strait is bordered by Greenland to the east and by Ellesmere Island to the west. It is essentially a flow-through system, consisting of deep channels and shallower basins (from north to south: Robeson Channel, Kennedy Channel, Kane Basin and Smith Sound; Fig 1). It is about 500 m deep, and its width varies from about 20 km across Robeson Channel at its northern end to around 150 km across Kane Basin [29]. From September to June, more than 80% of Nares Strait is covered by sea ice [30].

**Fig 1.**
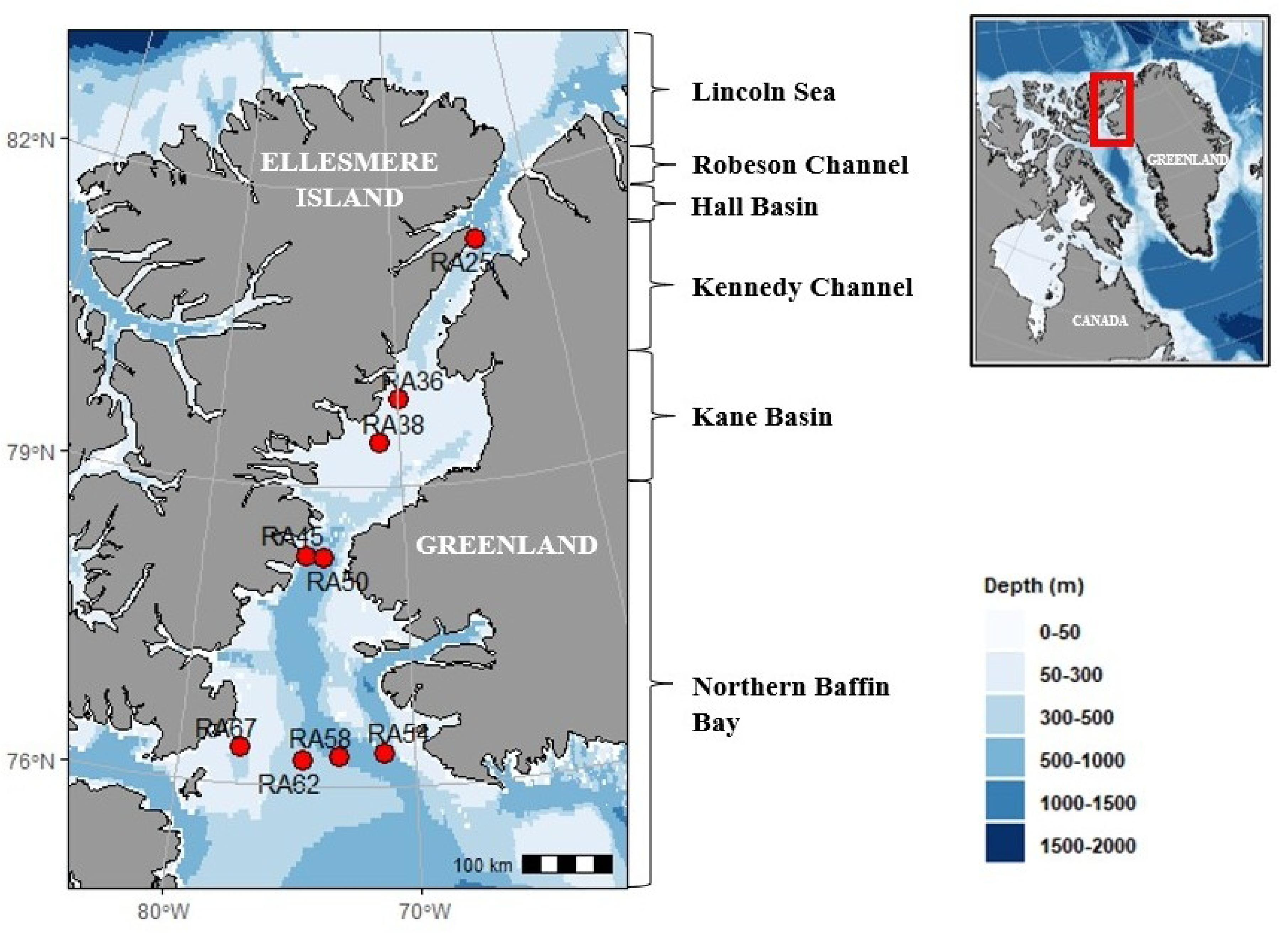
Location of stations sampled for sediment incubations in Nares Strait between Arctic Canada and Greenland in September 2024.

Nares Strait is affected by powerful southward winds, channelled by the steep terrain that rises to 2000 m above the sea surface on both sides of the strait. Currents also flow southwards from the Arctic Ocean through Nares Strait. This Arctic water, less salty than Atlantic water found in Baffin Bay, is composed of nutrient-rich Pacific-origin water and freshwater from melting sea ice and rivers. In addition to the liquid freshwater, Nares Strait seasonally exports an estimated 4-8 mSv of ice southwards [31]. The magnitude and the timing of sea ice export in this region largely depends on the ice arches that usually form at the northern and southern end of the strait and can remain stable for months at a time preventing mobile ice from flowing southward [32,33].

### Field sampling

Nine stations were sampled in September 2024 during the RefugeArctic cruise aboard the research icebreaker CCGS Amundsen along a latitudinal gradient in Nares Strait in the Canadian Arctic (Fig 1 and Table 1). Three stations, characterized by permanent landfast ice, were located in Kenedy Channel (RA25) and Kane Basin (RA36 and RA38). Two other stations with seasonal landfast sea ice coverage were located in Smith Sound (RA45 and RA50). The remaining four stations in the North Water Polynya are permanently free of ice (RA54, RA58, RA62 and RA67).

**Table 1.**
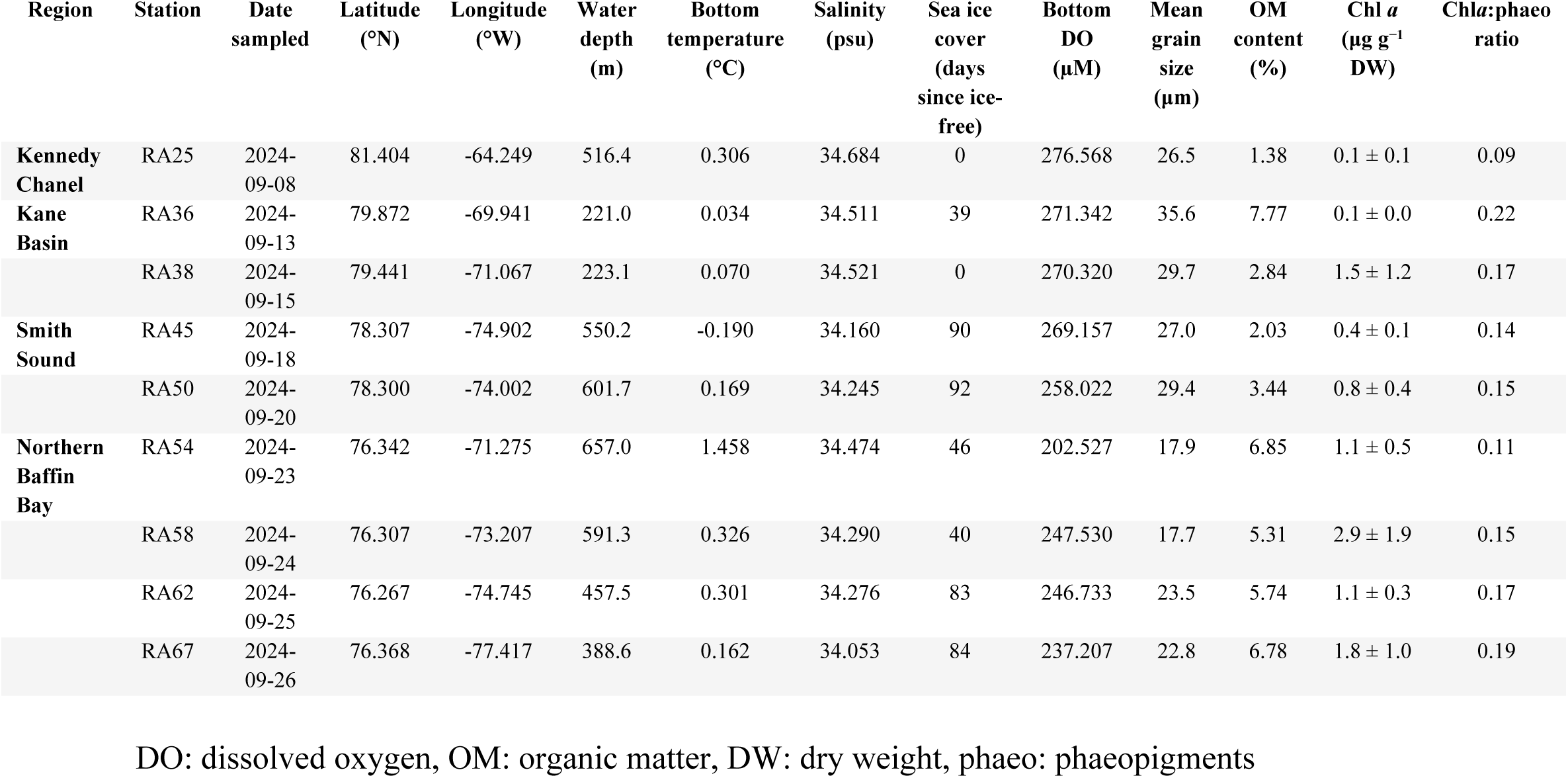
Station names of oxygen consumption measurements, sampling dates, locations in Nares Strait and environmental variables measured.

At each station, an USNEL Box corer (50 cm × 50 cm × 30 cm) was deployed at the seafloor for sediment collection (n=1 per station due to time limitations at station). From each box core, three sub-cores of 10 cm diameter and approximately 20 cm sediment depth were taken for assessing oxygen uptake and nutrient remineralization in incubations. In addition, the upper 1 centimetre of sediment was collected outside the sub-cores with 10 mL truncated syringes of an area of 1.5 cm^2^; n=1, for determining chlorophyll *a* (Chl *a*) and phaeopigment (phaeo) concentrations (mg m^−2^), organic matter content (% of dry sediment) and grain size (%). The samples for the food supply indicators (i.e., sedimentary Chl *a* and phaeopigments) were immediately frozen at −80 °C.

The samples for the organic matter and the grain size were frozen at −20 °C for latter analysis in the laboratory. The presence and extent of sea ice cover at the sampling sites were determined from the Canadian Ice Service (https://iceweb1.cis.ec.gc.ca/Archive/page1.xhtml). A window of 2 weeks before the sampling with a concentration of ice less than 20% (i.e., from open water to very open drift) was considered as ice-free.

### Incubations

Benthic remineralization function was measured as the combination of oxygen, nitrite, nitrate, phosphate, silicate and ammonium fluxes at the sediment-water interface. All fluxes were calculated in μmol m^-2^ d^-1^, except oxygen which was measured in mmol m^-2^ d^-1^. For this, shipboard incubations of sediments were performed as followed:

After retrieval and preparation on deck, sub-cores were pushed to ca. 15-20 cm into the sediment and taken immediately to a dark and temperature-controlled room (4 °C). Sediment cores were carefully topped with bottom water collected with the rosette at the same station. Incubation cores were acclimated for 6-8 h to allow suspended particles to settle to the sediment surface. During this period, the water was saturated with oxygen using an air aquarium pump to avoid suboxic conditions. After the acclimation period, air bubbles were carefully removed, and sediment cores were sealed with a lid equipped with a magnetic stirrer and a sealable hole for water sampling.

Sediment cores were incubated until a maximum of 20% of available oxygen was consumed to avoid suboxic conditions and biogeochemical transformations [34]. The incubation period was usually 24 to 48 h. Three supplementary incubation cores, incubated with bottom water only, served as controls to assess oxygen uptake and nutrient changes due to processes in the water column or due to sample handling.

### Benthic fluxes measurements

Oxygen flux was measured continuously with oxygen sensor spots SP-PSt3-NAU attached on the inner wall of each incubation cores. Measurements were made with a polymer optical fiber attached on the outside of the incubation cores (Fibox 4, PreSens, Regensburg, Germany).

To determine changes in nutrient concentrations (NH_4_ ^+^, NO_3_ ^−^, NO_2_ ^−^, SiO, PO ^3−^), water samples were collected at the beginning, midpoint and end of incubations (100%, 90% and 80% O_2_). The sampled water was replaced immediately by an equivalent volume of bottom water, with a known solute concentration. To prevent spurious measures, the volume of water withdrawn never exceeded 10% of the total volume of the incubation core. Water samples were filtered through 0.7 μm GF/F filters using acid-cleaned plastic syringes.

Measurements of nitrite, nitrate, phosphate and silicate were performed using a Bran and Luebbe Autoanalyzer 3 applying colorimetric methods adapted from Grasshoff et al. [35]. For ammonium measurements, water samples were analyzed onboard using a Turner Design fluorimeter following the method proposed by Holmes et al. [36]. The detection limit was 0.100 μmol L^−1^ for ammonium, 0.010 μmol L^−1^ for nitrate and nitrite, 0.020 μmol L^−1^ for phosphate and 0.016 μmol L^−1^ for silicate.

Oxygen and nutrient fluxes were assessed from the slope of the linear regression of oxygen and nutrient concentration versus time of incubation and corrected for the solute concentration in the replacement water measured at each filling [12,37,38]. Median flux determined in the control core was subtracted from each sediment core measurement. A positive nutrient flux indicates a release of nutrients from the sediment into the water column, while a negative flux signifies an uptake of nutrients from the water column into the sediment.

### Macrofaunal diversity

#### Taxonomic diversity

Following incubation, sediment cores were sieved through a 500 μm mesh sieve to collect infauna. Organisms were fixed with 4% formaldehyde solution. They were sorted under a dissecting microscope in the laboratory and identified to the lowest possible taxonomic level. We determined density (N; mean ± SE individuals) for each taxon, wet biomass (mean ± SE g) and taxonomic richness as the number of taxa present in each sediment core (mean ± SE). We also calculated the Shannon-Wiener (H’) and Pielou’s evenness (J’) diversity indices for each sediment core using the “vegan” R package [39]. Taxa not identified to the species level were distinguished from other specimens (e.g. sp. 1) and classified as morpho-species. Where such consistency across the study region was not achieved, specimens were grouped into the lowest common taxon.

#### Biological traits and functional diversity

Subsequently, species were classified into functional groups according to their traits in terms of feeding mode, body size, mobility and bioturbation influence (Table 2, Table S1). Traits were chosen based on their presumed influence on benthic remineralisation. Biological traits were collected for each taxon from the Arctic Traits Database and other published sources [8,12,40,41]. When biological traits information was unavailable for a specific taxon, we obtained information from one taxonomic rank higher. We used fuzzy coding that allowed more than one functional trait for a given taxon for each category and scored from 0 to 1 based on the extent to which they displayed each trait. For example, the cumacean *Eudorellopsis integra* can alternate between deposit and filter feeding depending on environmental conditions, so these two traits each scored 0.5 for the feeding type category.

**Table 2.**
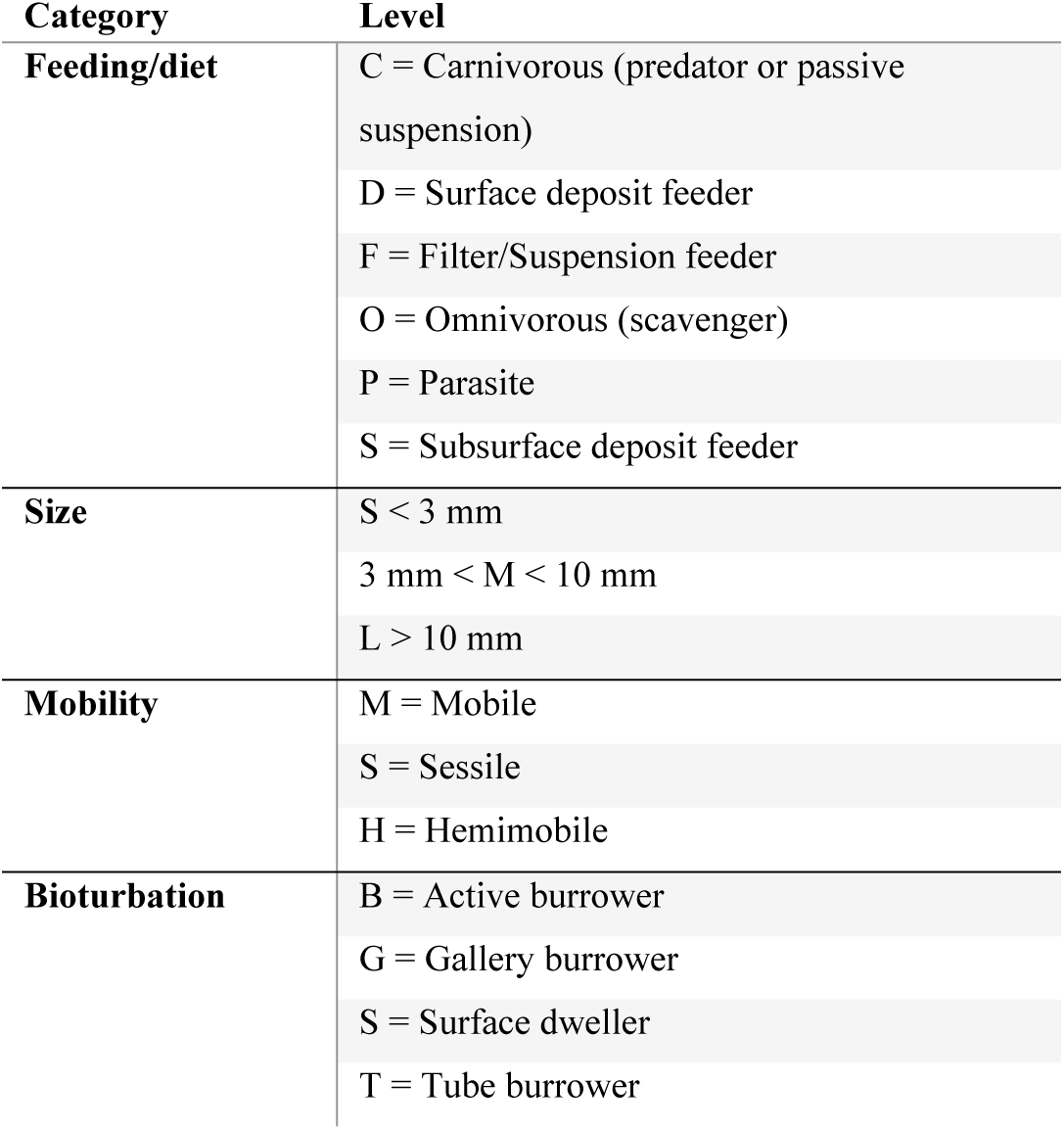
Categories of traits of Arctic benthic invertebrates and their levels (modalities) used to define functional groups.

Trait category scores for each taxon and taxa abundance matrices were used to obtain functional diversity (FD) indices using the “FD” package in R [17]. We then computed the following multidimensional FD indices for use in our analyses: functional richness (FRic), functional evenness (FEve) and functional divergence (FDiv) [17,18]. FRic represents the total functional space occupied by the community [18]. FEve, a unitless metric constrained between 0 and 1, describes the extent to which faunal abundances are evenly distributed within the trait space, where a value of 1 represents a perfectly even community [7]. FDiv quantifies heterogeneity of traits by representing the probability that 2 taxa picked at random from the community exhibit the same trait value [42].

In addition to the above metrics of functional diversity, we also determined community-level weighted means (CWMs) for each trait. CWM represents the probability that an individual organism drawn at random from the community will exhibit a given trait modality [7,43]. For example, a CWM of 0.6 for the trait modality ‘scavenger’ indicates a 60% probability that a random individual drawn from that community would be a scavenger.

#### Sediment properties

Chlorophyll *a* and phaeopigment concentrations were quantified using a modified protocol of Riaux-Gobin and Klein [44] and Link et al. [34]. We placed 2 g of wet sediment (the top centimeter of the sediment) in 10 mL of 90% acetone (v/v) for 24 h at 4 °C and then analyzed the supernatant before and after acidification (HCL 5%) using a TURNER Design 10AU fluorometer. Then, the sediment was dried and weighed to standardize pigment concentrations per gram of dry sediment. We sampled the top centimeter of the sediment sub-cores to quantify organic matter (OM) content by loss of ignition (500 °C). The water content was determined by comparing the mass of wet and dried sediment. We sampled the first five centimeters of sediment to determine granulometric properties (sediment mean grain size; MGS) with a Laser Diffraction Particle Size Analyzer LA-950 HORIBA. No sieving was performed prior to analysis because no large particles were present in our sediment samples.

### Statistical analysis

#### Spatial variation of benthic fluxes

We used a mixed-model PERMANOVA design to test for spatial differences in remineralization rates between stations. The resemblance matrices quantifying between-replicate similarities for all six standardized fluxes (O_2_ and five nutrients) were calculated using Euclidean distances performed with 9999 random permutations. Missing data points were replaced using the ‘missing’ function in PRIMER-E software. We verified homogeneity of multivariate dispersion using the PERMDISP routine [45]. We further analyzed significant terms in the full models using appropriate pair-wise comparisons. Non-metric Multidimensional Scaling (nMDS) plots were used to visualize the resemblance patterns. We completed PERMANOVA and PERMDISP analyses in PRIMER 6 with the PERMANOVA+ add-on [41].

#### Benthic biogeochemical flux drivers

Two separate distance-based redundancy analysis (dbRDA) were performed using PRIMER-E software using the stepwise distance-based linear models permutation test (DistLM) to identify which subset of diversity and environmental variables best predicted the multivariate variation of five benthic fluxes. We determined the model with drivers that best explained variation in benthic fluxes using a stepwise routine that employed 9999 permutations based on the AICc selection criterion. Draftsman’s plots of predictor variables indicated high correlation (r > 0.85) between five of our predictor variables (phaeopigment concentration, salinity, biomass and CWM surficial) and other predictors; we therefore excluded these variables from the analysis.

To correct for data skewness, we applied a square root transformation to the predictor variables temperature, abundance and dissolved oxygen. The response variables phosphate and ammonium required square root and cube root transformations, respectively. Nitrite required a natural log (ln) transformation. Prior to running the DistLM, we normalized flux, environmental and diversity data using the “normalise” function in PRIMER-E.

#### Variation partitioning of benthic fluxes with diversity variables

Variation partitioning was used to determine relative contributions of taxonomic and functional diversity indices to benthic flux variation. The method followed the principle used by Belley et al., [10]. Briefly, this was done by (1) performing a dbRDA of the flux on the taxonomic diversity data, (2) performing a dbRDA of the flux on the functional diversity data, (3) performing a dbRDA of the flux on the taxonomic and the functional data combined, (4) computing the adjusted R^2^ of the three dbRDAs, and (5) computing fractions of adjusted variation by subtraction.

## Results

### Spatial variation of individual benthic oxygen and nutrient fluxes

Oxygen, nitrite and ammonium were mostly taken up by the sediments, while nitrate, phosphate and silicate were mostly released, with a few exceptions in the western NOW at station RA67.Sediment oxygen demand varied between 0.7±0 and 4.4±0.7 mmol O_2_ m^−2^ d^−1^ with the lowest values at the northernmost station in Kennedy Chanel (RA25) and the highest values at the central Kane Basin and western NOW (RA38 and RA67; Fig 2). This spatial pattern was also evident in the release of phosphate (from 1.5±1.0 to 61.2±16.4 μmol PO_4_^3-^ m^−2^ d^−1^), silicate (from 183.1±24.3 to 4808.8±894.2 μmol SiO_2_ m^−2^ d^−1^) and the uptake of ammonium (from 12.9±7.4 to 106.3±150.9 μmol NH_4_^+^ m^−2^ d^−1^). The patterns for nitrite and nitrate fluxes were more variable. While nitrite was taken up by the sediments at all stations (from 0.9±1.3 to 5.5±1.9 μmol NO_2_^-^ m^−2^ d^−1^), it was released in RA67 (34.7±19.9 μmol NO_2_^-^ m^−2^ d^−1^). On the opposite, nitrate was taken up by the sediments in RA67 (78.8±60.6 μmol NO_3_ ^-^ m^−2^ d^−1^), while it was released in all the other stations (from 65.1±13.8 to 331.6±104.3 μmol NO_3_ ^-^ m^−2^ d^−1^).

**Figure 2.**
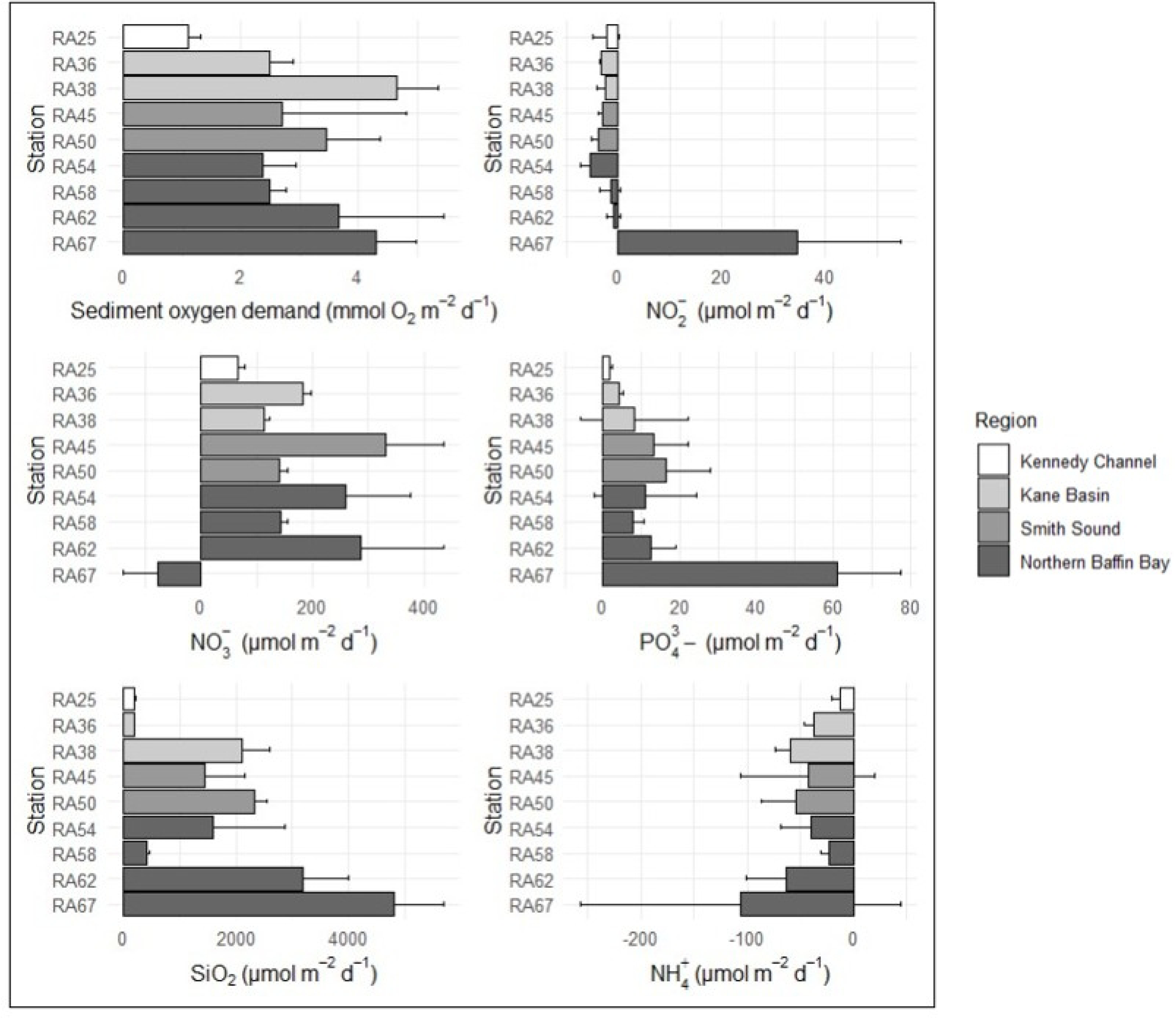
Mean (+ SE, n=3) of benthic flux of oxygen, nitrite, nitrate, phosphate, silicate and ammonium measured at each location. On the Y-axis are stations listed from south to north.

### Spatial variation in multivariate benthic fluxes

PERMANOVA indicated significant differences in benthic flux between stations and thus significantly greater variability in benthic fluxes among stations than within stations [P(perm) < 0.001]. The PERMDISP routine indicated no dispersion effect within our samples (p-value = 0.089). Pair-wise comparisons showed that benthic fluxes measured at RA67 differed significantly from fluxes measured at all the other stations [P(perm) < 0.05 in all 8 cases]. RA25 differed significantly from almost all stations except from RA54 [P(perm) = 0.095]. Moreover, the nMDS plot clearly showed that RA67 differed from the other stations, and that variability was greater across than within sites (Fig 3).

**Fig 3.**
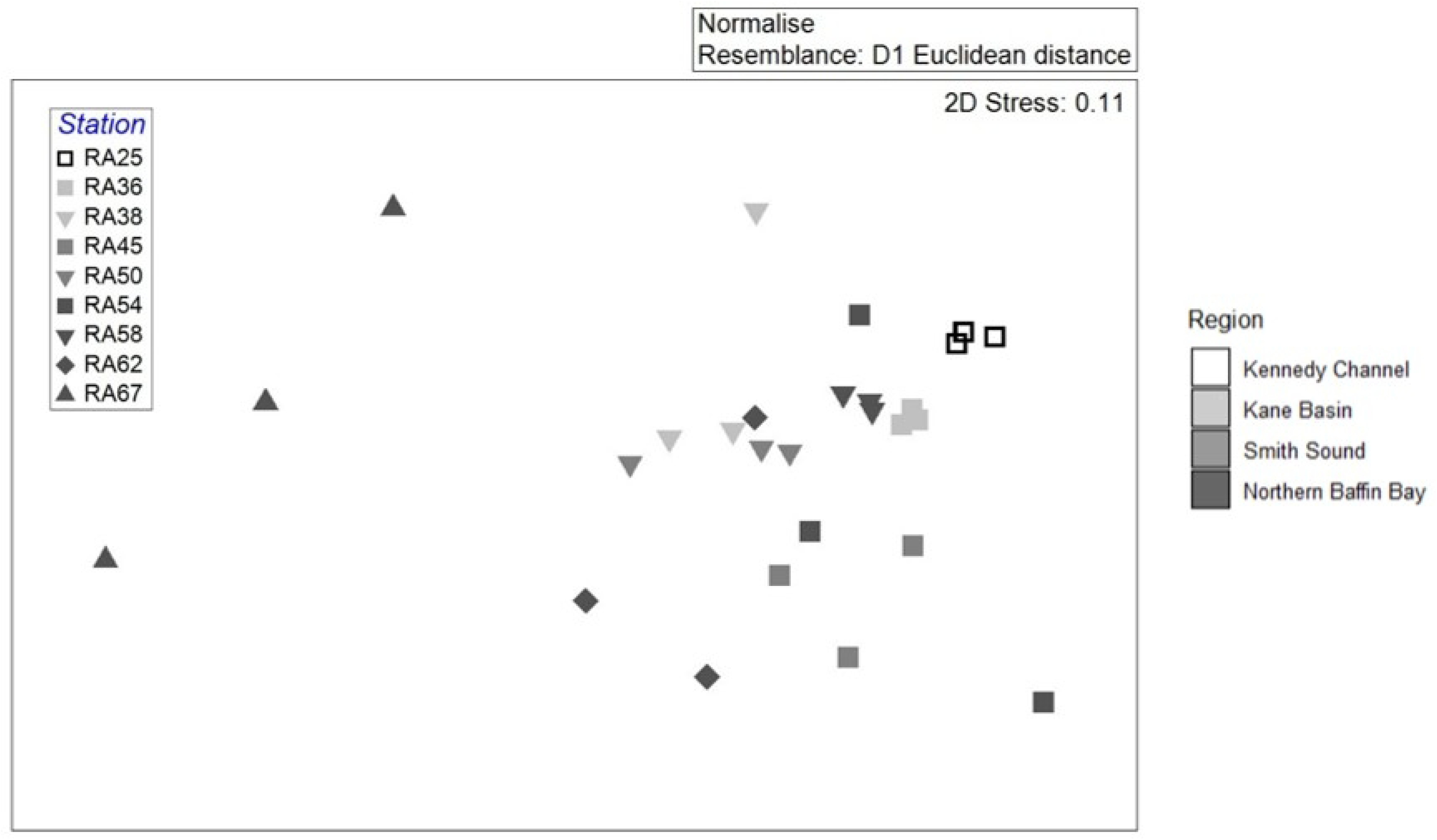
Non-metric multidimensional plot showing spatial pattern of benthic remineralization at each sampling station in Nares Strait in September 2024. The plot shows the relative distance of samples determined as Euclidean distance of the benthic biogeochemical fluxes. Each symbol represents one replicate.

### Environmental drivers of multivariate benthic fluxes

The best distance-based linear model (distLM) between benthic fluxes and environmental parameters explained 33.7% (R^2^ = 0.34, Adj. R^2^ = 0.25) of total benthic flux variation and included 3 variables (Fig 4A). Sea ice cover (number of days since ice-free) contributed most to the variation (13.9%), followed by mean grain size (MGS; 11.1%) and depth (8.7%) (Table S2).

**Fig 4.**
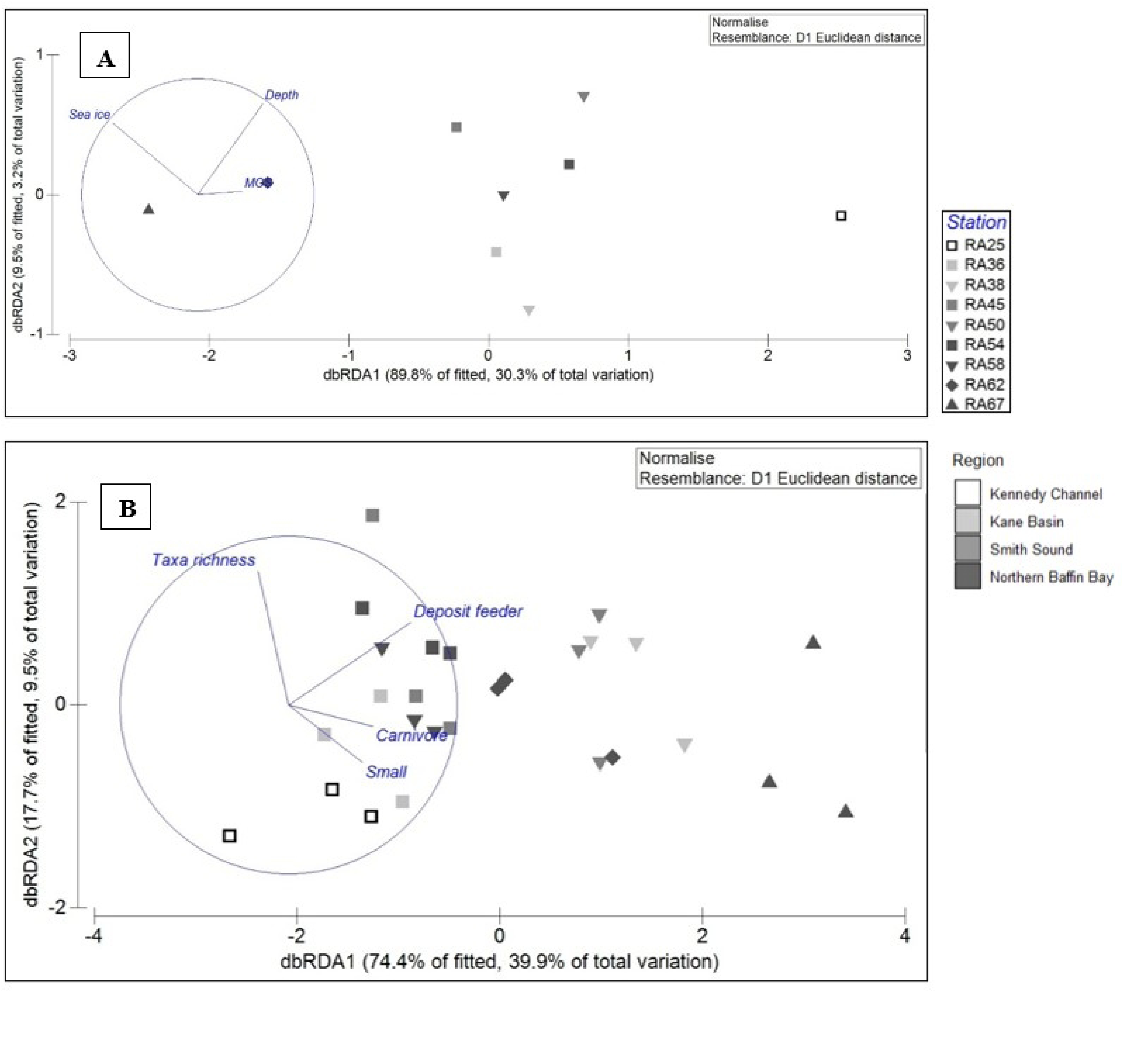
Distance-based redundancy analysis (dbRDA) plots of the distLM based on the environmental parameters (A) and the diversity indices (B) fitted to the variation in biogeochemical fluxes. MGS: mean grain size.

The first and the second axis account for 30.3 and 3.2% of the total flux variation, respectively. The most important parameters contributing to the first axis, explaining 89.8% of fitted flux variation, were sea ice cover and mean grain size (Fig 4A, Table S3). Depth was most strongly correlated with the second dbRDA axis, explaining 9.5% of the fitted flux variation.

### Macrobenthic diversity drivers of multivariate benthic fluxes

The best distLM model between benthic fluxes and macrobenthic taxonomic and functional diversity parameters explained 53.6% (R^2^ = 0.536, Adj. R^2^ = 0.452) of total benthic flux variation and included 4 variables. CWM of deposit feeders explained the most variation (22.6%), followed by taxa richness (12.1%), CWM of carnivores (11.5%) and CWM of small organisms (7.4%).

The first and the second axis account for 39.9 and 9.5% of the total flux variation, respectively. The most important parameters contributing to the first axis, explaining 74.4% of fitted flux variation, were CWM of deposit feeders, small organisms and carnivores, respectively (Fig 4B). Taxa richness was most strongly correlated with the second axis, explaining 17.7% of fitted flux variation.

### Benthic flux variation partitioning

Variation portioning of benthic fluxes between taxonomic and functional diversity indices indicated that taxonomic diversity indices alone explained 10.3% of total benthic flux variation, while functional diversity indices explained 40.4% (Fig 5, Table S4). Taxonomic and functional diversity indices shared 5.6% of the variation.

**Fig 5.**
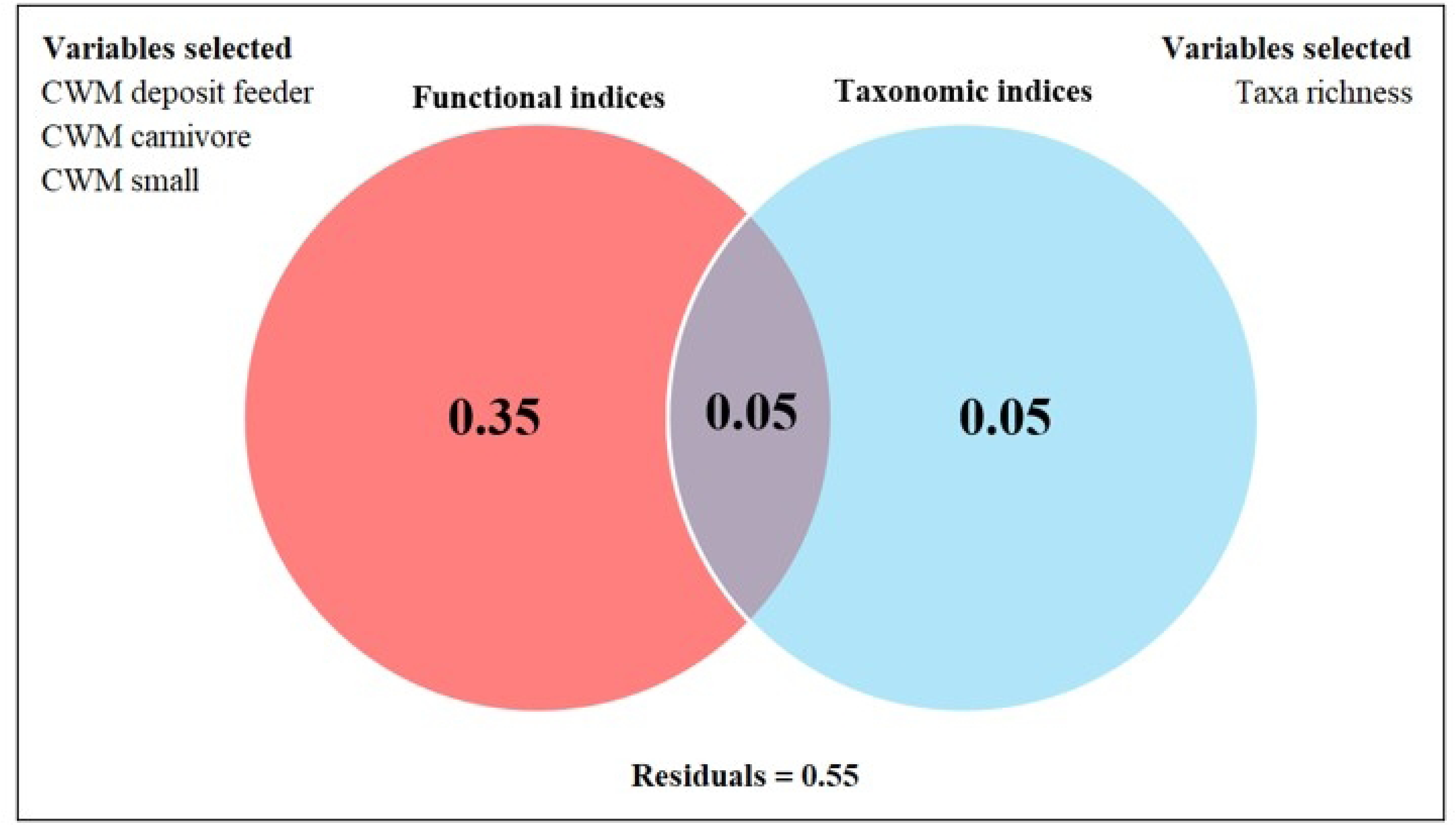
Venn diagram showing the variation partition of the functional and taxonomic diversity indices of macrobenthos in Nares Strait that contributes to explaining the benthic flux variation.

## Discussion

### Variability of oxygen and nutrient fluxes in Nares Strait

Overall, sediment oxygen demand measured in this study (0.7-4.4 mmol O_2_ m^−2^ d^−1^) exhibits similar values to those reported in previous Arctic studies on continental shelves around similar depths in summer/autumn. Comparable studies were conducted in the Baffin Bay and Canadian Arctic Archipelago (0.5-5.4 mmol O_2_ m^−2^ d^−1^; [12,47,48]), in the Beaufort Sea (0.5-11.5 mmol O_2_ m^−2^ d^−1^; [12,49,49]) and in the Barents Sea (2.3-7.3 mmol O_2_ ^-^ m^−2^ d^−1^; [37,50,51]).

However, the station in the Kenedy Channel (RA25) stood out through much lower sediment oxygen demand and nutrient fluxes. The sea ice conditions encountered at this station was more extensive and the ice much thicker, which prevented the research vessel from continuing its way North. Indeed, this region lies within the Last Ice Area, where the oldest and thickest sea ice in the Arctic Ocean is found [23,27,52]. The low SOD values at this station (0.7±0.0 mmol O_2_ m⁻² d⁻¹) were similar to values already recorded in deeper regions of the Arctic Ocean, such as the Central Arctic deep-sea basin (0.1-1.2 mmol O_2_ m^−2^ d^−1^; [53]), Fram Strait (0.5-1.9 mmol O_2_ m^−2^ d^−1^; [54]) and the deep regions of Laptev Sea (0.16-1.16 mmol O_2_ m^−2^ d^−1^; [51]). Although this station was located on the shelf and shallower than other stations (516m water depth), the permanent sea ice cover encountered probably limited primary production resulting in low export of organic matter to the seafloor and, consequently, benthic ecosystem functioning [56–59]. Indeed, the very low quantity of chlorophyll pigments found in the sediments of this station (0.1±0.1 μg g^−1^ DW Chl *a;* 1.1 ± 0.6 μg g^−1^ DW phaeopigments; Table 1) were very low and similar to the values found in the deeper regions of the Arctic.

In contrast to sediment oxygen demand, benthic nutrient fluxes (nitrite, nitrate, phosphate, silicate and ammonium) have only been measured on few occasions in the Canadian Arctic [8,12,49], Pacific Arctic [60,61] and isolated stations in Greenland [37,62–66]. Similar to the SOD measurements, nutrient values measured in our study were also within the range found in other Arctic regions [19]. However, an exception was identified in the western NOW. The benthic nutrient fluxes measured at RA67 differed significantly from those measured at all the other stations. This station, in western NOW at 389 m is in a region considered an advective sink for particles produced in the east of the polynya and subsequently transported by the net polynya circulation [29,67]. Consequently, sediment traps are increased in this region off Ellesmere Island, where particle accumulation and advection optimise conditions for sinking fluxes [68]. As a result, increased benthic oxygen demand has already been measured at this location and was suggested to coincide with the highest rates of carbon input measured by sediment traps and highest levels of sediment pigments [47]. However, our measurements are the first to record benthic nutrient fluxes in this area of the NOW and cannot be compared to previous studies.

Similarly heterogenous composition of benthic fluxes were observed in other regions of the Canadian Arctic [12] and were attributed to complex interactions with environmental [49,69] or faunal parameters [70,71]. However, none of the environmental or diversity variables measured at this station differed significantly from those at the other stations. Therefore, we suggest that the missing components of the micro- or meiofauna communities might explain the patterns found. Indeed, different types of bacteria can influence the rates and pathways of nutrient cycling (e.g. annamox and denitrification) in Arctic sediments [66,72,73]. Furthermore, growing evidence suggests that meiofauna can also influence these processes by stimulating the activity of the bacterial community in sediments [5,74,75]. It is therefore important that future studies consider all biotic compartments.

### Environmental drivers

Several environmental control factors for benthic remineralization have been suggested as underpinning the biogeochemical fluxes of carbon and nutrients in the Arctic [19,53,54]. Generally, labile carbon supply, temperature, sea ice cover, sediment type and water depth are the key factors controlling oxygen and nutrient fluxes in the Arctic. Many of these factors can vary spatially across regions. In particular, Nares Strait features multiple forms and ages of sea ice, from multiyear, first-year sea ice to summer sea ice-free. This region features a variety of sediment types and sizes due to the complex and strong currents in the strait, as well as glacially transported sediments due to the proximity of large glaciers and ice caps on both sides of the strait [76,77].

In this study, sea ice cover explained most of the variability in benthic fluxes, followed by depth and mean grain size, which is in line with previous studies in the Arctic [19,54]. Although sea ice cover does not directly affect benthic oxygen and nutrient fluxes, it is linked to them through a cascade of interdependencies across ecosystem components, such as primary production, food supply, and sediment properties [50]. Indeed, our results suggests that sea ice plays a central role in the structure of the environment and, by extension, in benthic fluxes. For instance, the lowest fluxes of oxygen and nutrients were found at the northernmost stations, which were still not ice-free at the time of the sampling, whereas the highest values were found in the NOW, which was completely ice-free at the time of the sampling. These differences in the remineralisation rates are not directly due to the ice itself, but rather to its effect on the environment i.e., in this case, the sediment size.

Indeed, the mean grain size influences benthic fluxes by altering sediment-water interactions, with finer sediments offering greater surface area for organic matter processing and bacterial activity, leading to higher oxygen and nutrient fluxes, which is consistent with our results [78]. Our study revealed that the grain size pattern in Nares Strait followed a latitudinal trend. The coarsest material, and associated lowest benthic fluxes, were found in Robeson Chanel and Kane Basin under lasting landfast sea ice, while the finest materials, and highest benthic fluxes, were found in the NOW.

Noteworthy is the lack of a clear relationship between pigment concentration and benthic fluxes. Although food availability is considered to be one of the most important factors in determining benthic remineralization rates in the Arctic [19], it was not included in our model as we originally hypothesized. We believe this is mainly due to the sampling period taking place at the very end of the productive period, just before winter i.e., sea ice in the strait was beginning to reform. Because of that, our measurements could represent only the “leftovers” of the season. Indeed, the low Chl *a*:phaeopigment ratios indicated old, processed phytodetritus, suggesting that the sampling occurred months after the input peak. The low input of fresh organic matter is reflected well in the low benthic activity, with oxygen uptake values in our study (0.7-4.4 mmol m^-2^ d^-1^) similar to those observed in the deep Arctic Fram Strait during the summer (0.5-5.1 mmol m^-2^ d^-1^; [54]).

It has been suggested that, in the case of low sediment pigment concentration and where rapid deposit feeding groups are present, functional community composition may bias food supply proxies [12]. Indeed, upon reaching the seafloor, phytodetritus can be consumed very rapidly by the benthos [37,79,80], and a pigment signal from a spring bloom in surface sediments and associated benthic fluxes can quickly fade. Deposit feeders can respond quickly to a pulse of fresh organic matter [81] and the community composition of all our stations, except the stations under lasting landfast sea ice, were dominated by deposit feeders (Table S5).

### Diversity drivers

Benthic communities also exert great influence on benthic carbon remineralization [10,19]. Macrobenthic community characteristics in terms of abundance, biomass and richness can explain an important part of the fluxes variation, especially in soft-bottom habitats [12,34,50,82]. In addition, previous studies have shown that species identity and differences in characteristics, such as feeding mode, can influence the intensity of bioturbation [83,84].

Indeed, based on bioturbation activity of sediment-dwelling invertebrate communities, Solan et al. [85] noted a clear separation in community composition across the Barents Sea polar front, suggesting that faunal activity is strongly moderated by seasonal variations in sea ice extent that influence food supply to the benthos. However, interpreting the bioturbation processes and the assignment of the macrofaunal species to the correct functional group can be challenging. Here, we considered these different processes independently in order to better understand their influence on benthic remineralisation rates.

Based on the taxonomic and functional indices selected, deposit feeders explained the most variation in benthic fluxes. While it is known that deposit feeders often dominate Arctic soft-sediment systems [86–88], the role they play in benthic carbon remineralization is less understood. Indeed, deposit feeders acquire food by swallowing large volumes of sediment [81], and as a result influence the rate of remineralization of carbon and nutrients through bioturbation. But more importantly, deposit feeders are known to assimilate ice algae efficiently [90–92]. The higher uptake of sympagic material by Arctic deposit feeders compared to other feeding strategies is well documented in many different Arctic areas, such as in the Pacific Arctic shelf [93], the Bering and Chukchi Seas [94], the Canadian Arctic Archipelago [95] or the Barents Sea [96]. More recently, the same phenomenon has been observed upon different megabenthic organisms belonging to several feeding strategies in Nares Strait [79]. This is usually attributed to the large, early season sedimentation pulse of fast-sinking ice algae aggregates on the seafloor, which occurs when zooplankton grazing is low [92].

Indeed, as sympagic production can contribute up to half of the total annual primary production in high Arctic marine systems [57,97,98], this gives them an advantage over other feeding strategies in specific regions where ice algae can provide a substantial food input, such as Nares Strait. A recent study revealed that sympagic production made up a significantly larger proportion of the organic matter deposited on the seafloor in Nares Strait than pelagic production in late summer [79], and thus might indirectly explain, via food preference, why most stations were dominated by deposit feeders (Table S5).

Our redundancy analysis also highlighted the importance of other functional traits related to reworking the sediment matrix. For example, scavengers and predators contribute to bioturbation by physically disturbing sediments through their activities, such as ingesting and moving food remains, which can alter sediment structure and influence biogeochemical processes in an ecosystem. Together, these results highlight the importance of traits related to the biological mixing of sediments during feeding. For instance, in the Strait of Georgia, Belley et al. [10] identified funnel feeding as a key function provided by a small number of species and individuals of macrofauna which can affect benthic flux rates disproportionately relative to their abundance.

Surprisingly, functional richness (FRic) had a low explanatory power in the benthic flux variations relative to taxa richness in our study. Our results are in accordance with Link et al. [12] who found higher contribution of abundance and taxonomic diversity to benthic flux variations compared to functional richness in the North Water Polynya. Nonetheless, they highlighted that the abundance of gallery-burrowing polychaete species, *Lumbrineris tetraura*, also a deposit feeder, was an explanatory variable for the variations in the fluxes. Previous studies noted that in environments characterized by low functional richness, species richness can be more important in controlling benthic ecosystem function [99]. Indeed, in environments with low functional diversity, as is the case in our study, higher species richness can become more important than functional richness for ecosystem function, as multiple species may perform the same function, increasing functional redundancy. From an ecological point of view, this redundancy allows the ecosystem to maintain function even if some species are lost, which is especially important in environments with limited resources and where specific functional groups are crucial for processing organic matter, which appears to be the case in Nares Strait, especially at the northernmost stations.

Our variation partitioning results demonstrate that, overall, functional indices are still better predictors in benthic fluxes than taxonomic indices but that considering both together allows to explain more variation, especially in resource-limited environments, as is the case in Nares Strait.

## Conclusion

Our study indicates strong spatial variation in benthic fluxes in Nares Strait, largely controlled by variations in sea ice cover and feeding type composition of benthic fauna. From a biogeochemical point of view, we were able to discern two distinct regions: The northern part (Kennedy Channel) covered by multi-year ice, was distinct from the seasonally ice-covered areas (Kane Basin and the North Water Polynya). The very low benthic fluxes measured in the Kennedy Channel were as low as those in deeper regions of the Central Arctic Ocean basins. Given the climate crisis is a carbon crisis, the shift from perennial (multi-year) to seasonal sea ice, which is already occurring in Nares Strait and many other Arctic regions, also implies changes in the capacity of these areas to act as carbon sinks. Our study is the first to reveal the links between macrobenthic functional diversity and benthic fluxes in Arctic sediments. Our findings provide crucial support for assessing the ability of the Arctic benthos to process carbon in response to climate change. Future studies should conduct larger-scale and longer time-series monitoring surveys to enhance our ability to predict the effects of environmental changes on the functional structure of Arctic benthic communities.

## Acknowledgements

We would like to thank the CCGS Amundsen officers and crew, scientists and technicians for their support on board. Special thanks to G. Blais and S. Pagé for the help on board. We are also grateful to J. Tréau de Coeli for species identification, J. Gagnon for nutrient analyses and R. Strassburg-Huot for the help with pigment analyses.

## Supporting information

**Table S1. List of taxa identified in sediment cores and the accorded functional traits.**

**Table S2. Redundancy analysis of benthic fluxes against environmental drivers in Nares Strait in 2024.**

**Table S3. Redundancy analysis of benthic fluxes against diversity indices in Nares Strait in 2024.**

**Table S4. Results of the variation partitioning of the benthic fluxes between taxonomic and functional diversity.**

**Table S5. Community-level weighted means (CWMs) of each functional traits measured from the communities in the incubation cores.**

## Notes

### Competing Interest Statement

The authors have declared no competing interest.

